# Endoplasmic Reticulum membranes are continuously required to maintain mitotic spindle size and forces

**DOI:** 10.1101/2022.05.14.491942

**Authors:** Margarida Araújo, Alexandra Tavares, Diana V. Vieira, Ivo A. Telley, Raquel A. Oliveira

## Abstract

Membrane organelle function, localization, and proper partitioning upon cell division depend on interactions with the cytoskeleton. Whether, reciprocally, membrane organelles also impact on the function of cytoskeletal elements remains less clear. Here, we show that acute disruption of the Endoplasmic Reticulum (ER) around spindle poles affects mitotic spindle size and function in *Drosophila* syncytial embryos. Acute ER disruption was achieved through the inhibition of ER membrane fusion by the dominant-negative cytoplasmic domain of Atlastin. We reveal that when the ER is disrupted specifically at metaphase, mitotic spindles become smaller, despite no significant changes in microtubule dynamics. These smaller spindles are still able to mediate sister chromatid separation, yet with decreased velocity. Furthermore, by inducing mitotic exit, we found that nuclear separation and distribution are affected upon ER disruption. Our results suggest that ER integrity around spindle poles is crucial for the maintenance of mitotic spindle shape and pulling forces. Additionally, ER integrity also ensures nuclear spacing during syncytial divisions.

## Introduction

Cell division is often simplified to an isolated process of chromosome segregation by the mitotic spindle (Vitre et al. 2014) followed by scission of the cell membrane during cytokinesis (Mierzwa and Gerlich 2014). Thereby, the spindle apparatus assembles, attaches chromosomes, aligns them, and generates the force required to pull sister chromatids apart (Petry 2016; Maiato et al. 2017). However, in compartmentalized eukaryotic cells various organelles undergo extensive reorganization and distribution to the daughter cells during mitosis (Champion et al. 2017; Carlton et al. 2020). One of those organelles is the Endoplasmic Reticulum (ER), a tubular network that forms a continuum with the nuclear envelope (Goyal and Blackstone 2013). When the nuclear envelope breaks down in mitosis, allowing spindle microtubules (MTs) to interact with the chromosomes, the ER reorganizes in the vicinity of the spindle (Bobinnec et al. 2003). ER membranes have recently been proposed to hinder efficient chromosome segregation. When ER membrane biogenesis is increased by enhanced fatty acid synthesis, this leads to higher viscosity of the cytoplasm and ultimately to chromosome mis-segregation (Merta et al. 2021). Moreover, chromosomes that become unsheathed by the ER are more prone to segregation errors (Ferrandiz et al. 2022). Hence, ER reorganization may be a passive event, required for faithful mitosis and even distribution of this organelle to daughter cells (Smyth et al. 2015). Reciprocally, it has also been proposed that ER reorganization plays a functional role during mitosis. On one hand, the ER enclosing the spindle could act as a molecular exclusion barrier sorting or concentrating cell cycle and spindle relevant proteins (Schweizer et al. 2015). On the other hand, it is conceivable that the ER plays a mechanical role and balances spindle forces, thus adjusting e.g. the spindle length (Dumont and Mitchison 2009). This is mostly supported by the observation that functional disruption, at mitotic entry, of another membranous structure – the nuclear envelope – perturbs spindle assembly (Tsai et al. 2006; Liu and Zheng 2009; Ma et al. 2009; Civelekoglu-Scholey et al. 2010). At the molecular level, REEP proteins were shown to be involved in the exclusion of ER membranes from the spindle region during mitosis (Schlaitz et al. 2013). Furthermore, the ER targeting kinase TAOK2 is important for tethering of ER membranes to the MT cytoskeleton and for ER mobility along MTs during mitosis (Nourbakhsh, Ferrecccio, and Yadav 2021). Altogether, these studies exposed the role of membranes during mitosis and demonstrated that the association between the ER and MTs is important for spindle assembly. Whether this association is required continuously, even after unperturbed spindle assembly, is currently unknown.

Deciphering an actively contributing versus passively hindering role of the ER during mitosis is critical. However, investigating these potential roles is technically challenging due to the difficulty to perturb such a critical structure for cell physiology in a fast and temporally controlled manner. Here, we used microinjection approaches to disrupt acutely ER membranes in a metaphase-arrested state and examine mitotic spindle morphology and function upon loss of ER integrity. We uncovered that ER membranes surrounding the spindle pole are important for the maintenance of mitotic spindle architecture and forces.

## Results

To visualize the ER during *Drosophila* syncytial embryonic divisions, we performed live imaging and followed ER and nuclear envelope localization throughout the cell cycle. We generated flies expressing the chromatin marker H2B–mRFP1, and either the ER marker EYFP–KDEL or the ER membrane-shaping protein Reticulon-like protein 1 (Rtnl1) fused to GFP (GFP–Rtnl1). We also marked microtubules by injecting Alexa647-labelled tubulin (Fig. 1, Suppl. Video 1). KDEL is a small peptide sequence that targets proteins to the ER lumen (Frescas et al. 2006). EYFP fused to the KDEL sequence reports all ER, while Rtnl1 specifically reports locations of membrane reshaping. Consistent with this notion, we observed an extended KDEL-labelled ER network in the entire cortex of the syncytial embryo and throughout mitosis (Fig. 1A), with partial overlap with the nuclear envelope (Lamin–GFP, see Fig. S1, t=00:00). This suggests an interaction between these two membranous structures during interphase. However, at mitotic entry upon nuclear envelope breakdown, the concentration of ER in the vicinity of the spindle rose (Fig. 1A, Metaphase). The signal of GFP–Rtnl1 also increased, exclusively around the spindle, and most prominently at the spindle poles (Fig. 1B). This is consistent with prior reports showing an increase in ER proteins at spindle poles upon mitotic entry (Bobinnec et al. 2003; Diaz et al. 2019). However, it remains unclear how these changes in ER organization impart on ER morphology across different stages of spindle assembly. Of note, both ER reporters shown are excluded from the spindle and resemble an envelope (Fig. 1A-B, insets). At NEBD, we measured a reduction in the ER exclusion area (Fig. 1C) supporting a transient contraction of the ER at this stage. This reduction is not accompanied by a decrease in the perimeter of the ER envelope (Fig. 1D), due to an indentation of the ER envelope at spindle poles (Fig. 1C, inset). During metaphase and anaphase, the area and perimeter of the ER exclusion gradually increase, matching closely that of the spindle main body, disregarding the spindle poles (Fig. 1D). This comparative measurement emphasizes the shape similarity of the ER envelope and the spindle body. At telophase, the ER undergoes shape changes that accompanied nuclear envelope reformation (Fig. 1A-B, Telophase and Fig. S1). We also observed that both ER reporter proteins localized at the spindle midbody as previously reported (Bobinnec et al. 2003), suggesting that a considerable membrane reorganization occurs at this site (Fig. 1A-B, Telophase, arrow).

**Figure 1:**
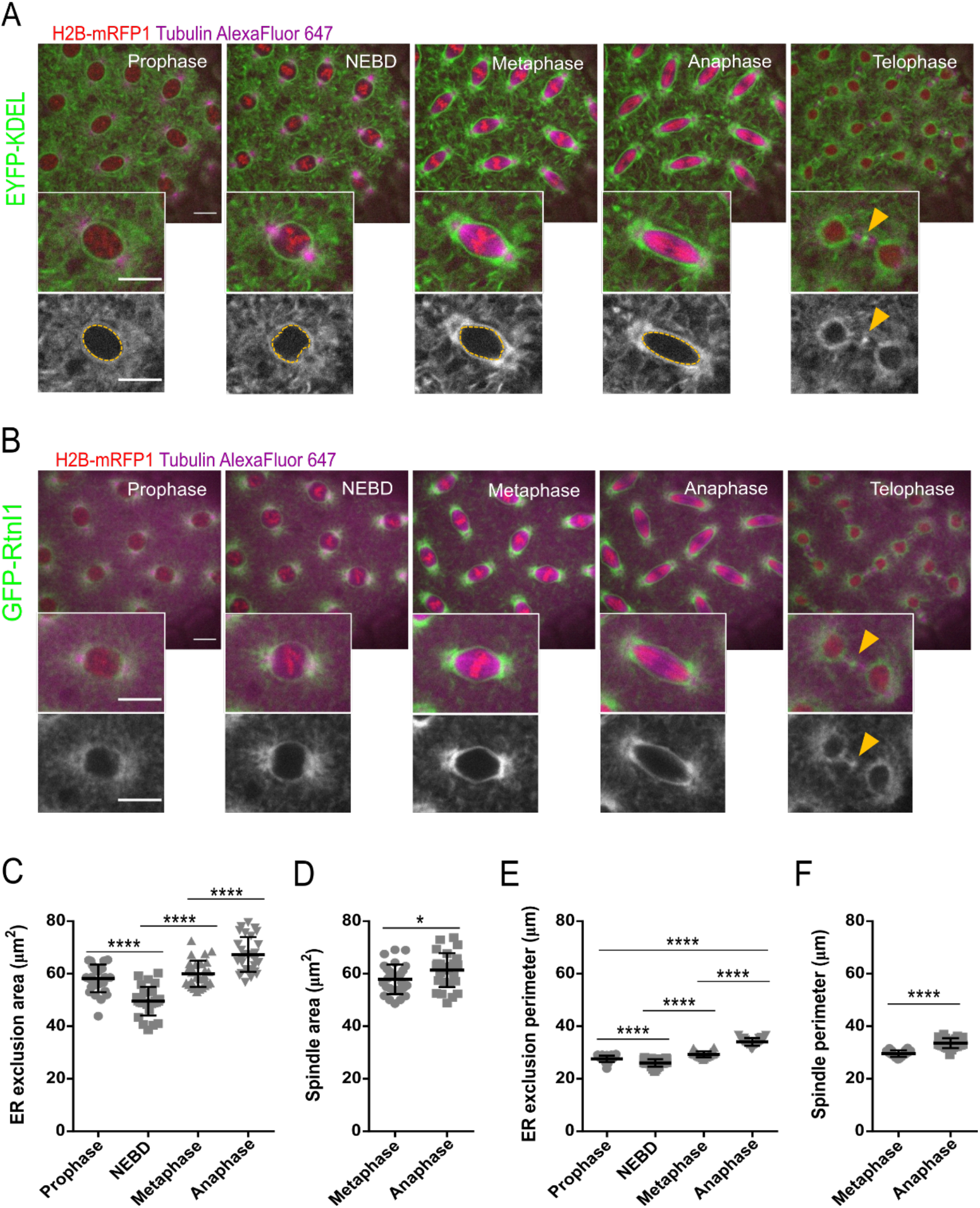
The ER forms an envelope surrounding the mitotic spindle in syncytial embryos. **(A-B)** Stills of embryonic divisions monitoring ER dynamics at different mitotic stages. ER was visualized with either EYFP-tagged ER retention sequence (EYFP–KDEL, green, A) or the ER-shaping protein Reticulon-like protein 1 (GFP–Rtnl1, green, B). Chromatin is labelled with Histone H2B–mRFP1 (red) and spindle microtubules with microinjected porcine Tubulin labeled with Alexa Fluor 647 (magenta). Grey panels depict ER labelling alone. Scale bar is 10 µm. Arrowheads show events of ER abscission at telophase. **(C-F)** Quantifications of the ER exclusion area (C), spindle area (D), ER exclusion perimeter (E) and spindle perimeter (F), measured at the middle plane of the nuclei using the EYFP–KDEL strain; sample size: N=7 embryos per condition, n=5 nuclei per embryo; statistical analysis was performed using one-way ANOVA, multiple comparisons, p value adjusted to multiple comparisons (Tukey) (C and E) or two-sided unpaired t-test (D and F); *p<0.05,****p<0.0001, n.s. = non-significant, p>0.05.

### ER reorganization throughout mitosis

Next, we wanted to understand if the observed ER shape changes, which closely follow those of the spindle body, arise from dynamic changes of ER membranes themselves. To this end, we performed Fluorescence Recovery after Photobleaching (FRAP) experiments using the GFP-tagged transmembrane protein Rtnl1, which labels membranes that are being re-shaped and tubular ER is being formed (Espadas et al. 2019). To circumvent the inherent changes in intensity and morphology during the cell cycle, which would impede proper FRAP analysis in these fast cycles, we have performed all the experiments in artificially arrested embryos. We first monitored the ER dynamics in interphase, by preventing mitotic entry with ectopic addition of the Cdk inhibitor p27 (Oliveira et al. 2010). Upon p27 injection, we observed that the ER maintained an interphase-like localization (Fig. 2A). For FRAP studies, we bleached two different arrested nuclei in distinct regions (ROIs) and imaged their fluorescence recovery over time (Fig. 2A, red circles, Suppl. Video 2). We observed a high turnover of ER membranes, with half-times of recovery in the order of seconds. ER membranes localized proximal to the spindle poles are more dynamic compared to those localized at the equator (Fig. 2A, t_1/2_ poles: 11.4 ± 6.2 s; t_1/2_ equator: 21.0 ± 8.6 s). However, the GFP–Rtnl1 mobile fraction at the poles is lower compared to the virtually complete recovery at the equator in interphase-arrested embryos (poles: 0.81 ± 0.06; equator: 0.9 ± 0.08). Our observations are in contrast to previous studies in yeast and mammalian cells, which detected a much larger immobile fraction for Rtn1/Rtn4a (Shibata et al. 2008). These findings suggest a high level of reshaping of ER membranes in *Drosophila* embryos.

**Figure 2:**
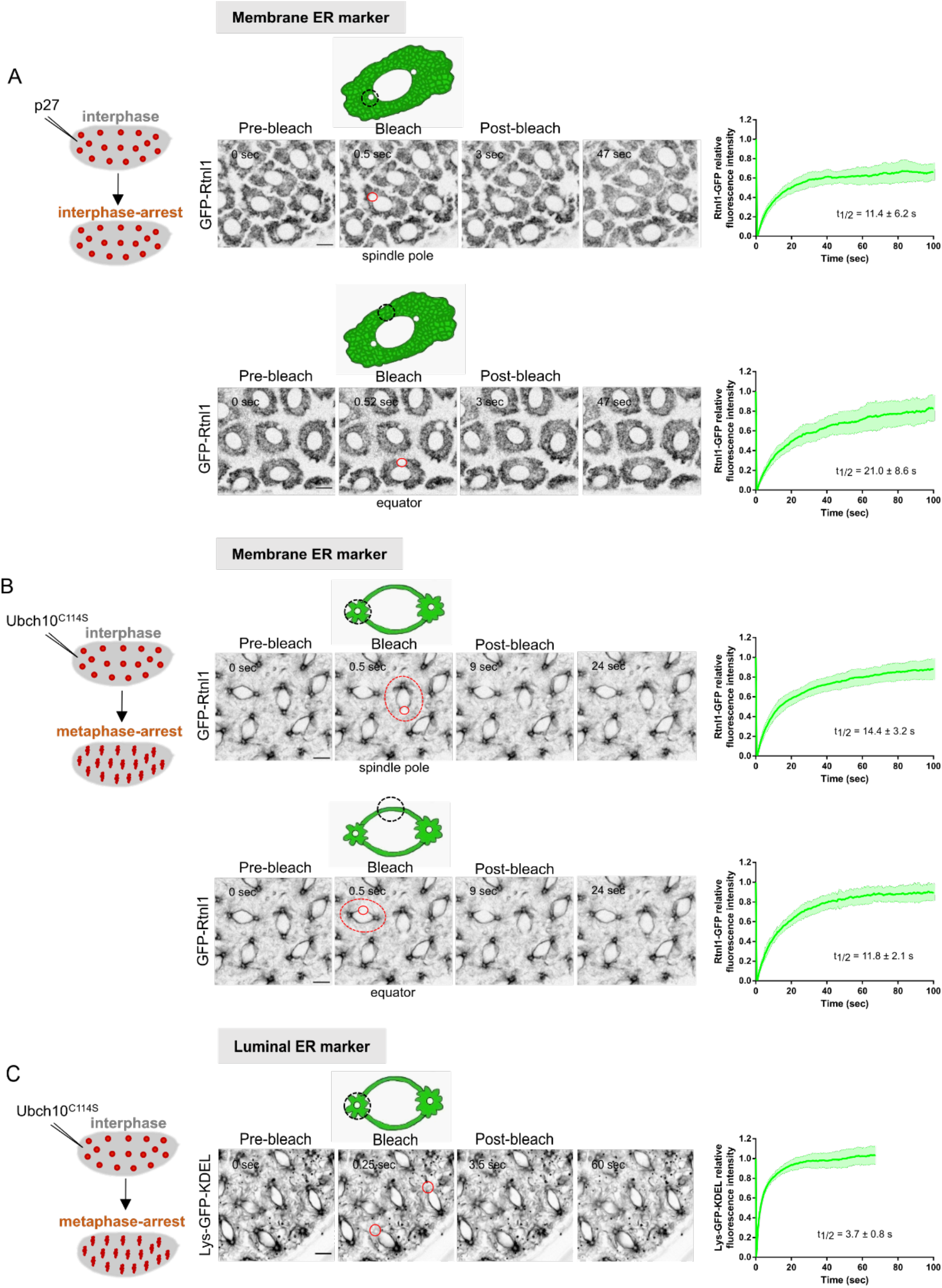
ER dynamics in interphase- and metaphase-arrested nuclei. **(A)** Fluorescence recovery after photobleaching (FRAP) of the membrane ER marker GFP– Rtnl1, in embryos microinjected with the Cdk inhibitor p27 to induce an interphase-arrest. Recovery of Rtnl1–GFP fluorescence intensity was quantified upon bleaching of the ER at the spindle pole region (upper panel; N=6 embryos, n=12 nuclei) or at the equator (lower panel; N=5 embryos, n=9 nuclei). **(B)** FRAP analysis of GFP–Rtnl1 in embryos microinjected with the dominant-negative version of the E2 ubiquitin-ligase UbcH10 (UbcH10^C114S^) to induce a metaphase-arrest. Graphs depict recovery of Rtnl1–GFP intensity upon bleaching at the spindle pole region (upper panel, N=9 embryos, n=17 nuclei) or at the equator (bottom panel, N=9 embryos, n=15 nuclei); red circles depict the area bleached in each experimental condition. **(C)** FRAP analysis of Lys–GFP–KDEL in metaphase-arrested embryos (UbcH10^C114S^-injected). Graph depicts recovery of Lys– GFP–KDEL after photobleaching at spindle poles (N=5 embryos, n=10 nuclei). In all experiments, average half-times of recovery are depicted, presented as mean ± standard deviation. Scale bars are 10 µm.

Changes in ER morphology are coupled to the cell cycle and are dependent on Cyclin A activity during *Drosophila* embryonic nuclear divisions (Diaz et al. 2019). To address if this morphological re-organization is accompanied by a change in the dynamic behavior of ER membranes, we repeated the same analysis in embryos arrested in metaphase. For this, embryos were microinjected with UbcH10^C114S^, a dominant-negative catalytically dead version of the E2 ubiquitin-conjugating enzyme necessary for anaphase onset (Oliveira et al. 2010). We bleached two different nuclei in distinct regions of interest (ROIs) and imaged their fluorescence recovery over time (Fig. 2B, red circles). Signal recovery of the ER shaping protein Rtnl1 was in the same order of magnitude (Fig. 2B, Suppl. Video 3). Upon metaphase-arrest, we detected a slightly slower recovery of Rtnl1– GFP intensity at spindle poles compared to the equatorial region (poles: 14.4 ± 3.2 s; equator: 11.8 ± 2.1 s). Interestingly, we observed virtually complete recovery of intensity in both cases (mobile fractions at pole: 0.92 ± 0.06; equator: 0.95 ± 0.04). This signal dynamics observed is slightly lower, but within the same magnitude of the turnover observed for the luminal ER reporter Lys–GFP–KDEL (Fig. 2C, t_1/2_: 3.81 ± 0.46 s, Suppl. Video 4), which reflects ER shape changes occurring in the time scale of free diffusion events inside the ER. This is in agreement with previous studies, where rapid recovery of intensity was reported for both *Drosophila* oocyte fusome and syncytial embryos expressing Lys–GFP–KDEL (Snapp et al. 2004; Frescas et al. 2006). These findings suggest that the ER in mitosis is continuously undergoing significant reorganization.

### Ectopic addition of cytATL changes mitotic ER topology at spindle poles

To explore how these dynamic ER membranes could impact on the overall architecture of mitosis, we developed a strategy to acutely perturb the ER in metaphase-arrested embryos and follow the consequences in real-time. To this end, we made use of the cytosolic domain of the ER membrane fusing protein Atlastin. This truncated version has been used as a dominant-negative reagent in *Xenopus* egg extracts to impair the membrane fusion activity of the native form on the ER (Wang et al. 2013; Wang et al. 2016; Kutay et al. 2021). *D. melanogaster* cytATL was expressed in and purified from bacteria (Suppl. Fig. 2) and microinjected into syncytial embryos. To monitor the ER specifically during metaphase, embryos were previously arrested with UbcH10^C114S^, as above (Fig. 3A). We observed that subsequent microinjection of cytATL in metaphase-arrested embryos alters mitotic ER topology compared to controls (Fig. 3B-C, Suppl. Video 5). Over time, EYFP–KDEL labelled ER membranes lose their linear arrangement, typical of a tubular structure, and a acquire diffuse and homogeneous appearance (Fig. B-C, t=10:00 min). Quantitative analysis reveals that there is no change in the mean intensity of the ER reporter at spindle poles, with or without cytATL injection (Fig. 3D). However, the spatial distribution of the signal altered, as evidenced by the significantly decreasing variance upon cytATL injection (Fig. 3E). This suggests that the ER contents remain but their distribution or local concentration changes. In contrast, centrosome-distal regions do not change significantly both in mean and distribution of the signal (Fig. 3D-E). These findings reveal that the disruptive effect of cytATL is exclusively observed at ER membranes surrounding spindle poles. In addition to the change in spatial ER concentration, ectopic addition of cytATL caused a significant reduction in the ER exclusion zone (Fig. 3F). We therefore concluded that ER membranes at the spindle poles are more sensitive to the disruptive effect of cytATL, whose addition leads to acute changes in morphology at this sub-region of the ER network.

**Figure 3:**
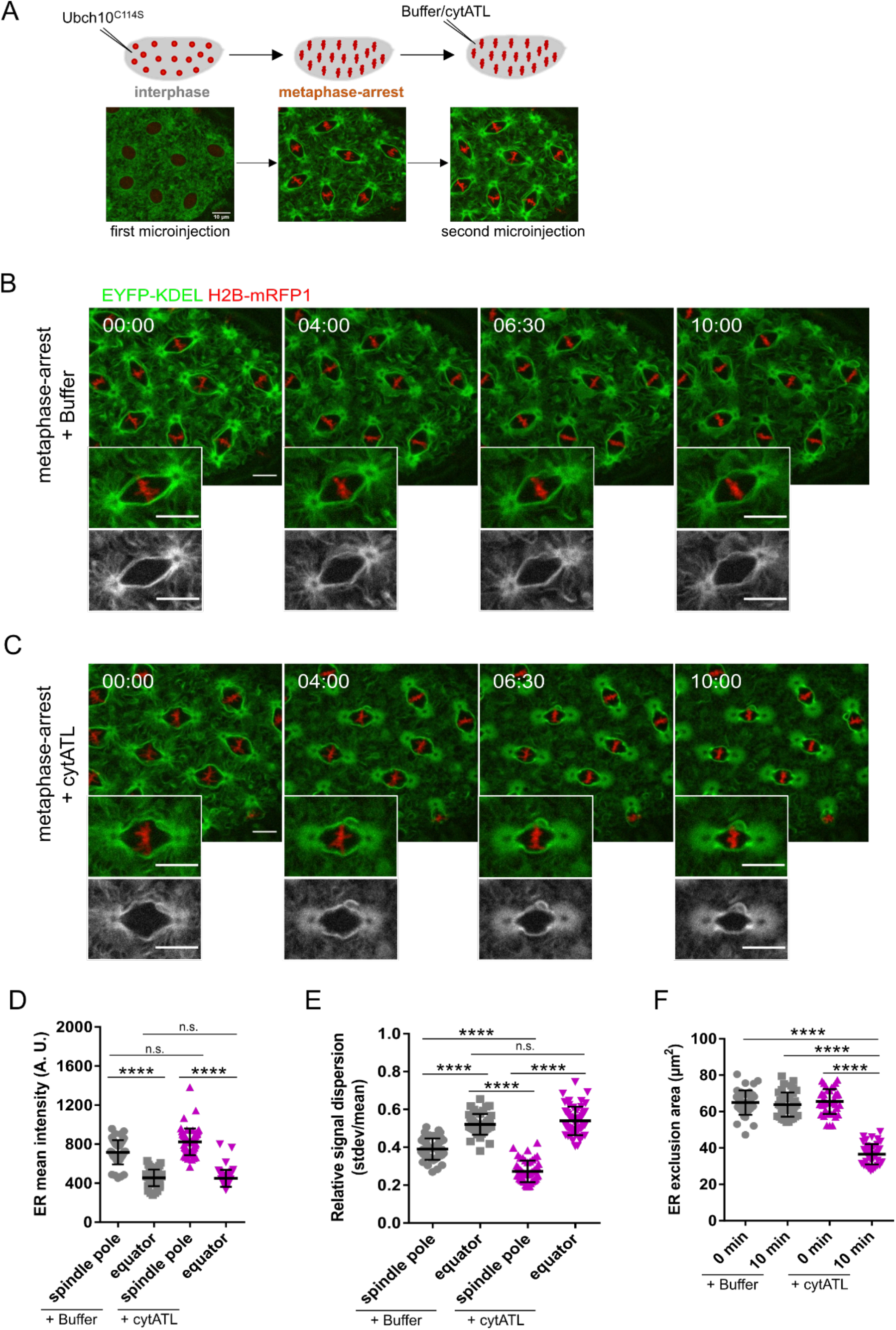
Acute ER disruption is achieved by ectopic addition of cytATL. **(A)** Experimental layout for acute ER disruption: embryos were arrested in metaphase by microinjection with UbcH10^C114S^ and allowed to reach chromosome alignment. After arrest (∼5 min) embryos were subjected to a second microinjection. ER dynamics was monitored by EYFP–KDEL (green) and chromatin by H2B–mRFP1 (red). **(B-C)** Stills depicting changes in ER morphology (EYFP–KDEL, green) after microinjection of buffer (B) or the dominant-negative cytATL protein (C) in metaphase-arrested embryos; time (min:s) is relative to the second microinjection (buffer/ cytATL). Grey panels depict ER channel alone. Scale bar is 10 µm. **(D)** Quantitative analysis of mean fluorescence intensity of EYFP–KDEL in spindle poles or equatorial regions 10 min after buffer/cytATL injection. **(E)** Coefficient of variation, calculated by the ratio of the standard deviation over the mean (stdev/mean), 10 min after buffer/cytATL injection. **(F)** ER exclusion area in control (buffer-injected, grey) and cytATL (magenta) conditions at the first (t=0 min) and last (t=10 min) time point of the time-lapse. Statistical analysis using N=10 embryos, n=5 nuclei per embryo. Asterisks represent statistical significance derived from One-way ANOVA, multiple comparisons, p value adjusted to multiple comparisons (Tukey). **** = p<0.0001, n.s. = non-significant, p>0.05.

### Acute disruption of spindle pole-proximal ER membranes decreases mitotic spindle size

Having established a method that acutely disrupts ER integrity in mitosis, particularly at centrosome-proximal regions, we next sought out to evaluate the effect this disruption has on mitotic spindle architecture. For this, we co-injected UbcH10^C114S^ with porcine Tubulin labelled with AlexaFluor 647 to visualize spindle microtubules during the metaphase arrest (Suppl. Video 6). Upon cytATL-induced ER disruption, we found that the morphology of the spindle is significantly altered (Fig. 4B-C, t=10:00 min). Changes in spindle architecture are evidenced by a reduction in spindle length and width when compared to control embryos (Fig. 4D-E). Thus, we conclude that ER morphological changes mediated by the dominant-negative effect of cytATL lead to smaller mitotic spindle size. In addition to this effect, we also observed, at considerable frequency, the detachment of the spindle pole microtubule-organizing center (MTOC) upon ectopic addition of cytATL. To quantify this, we measured the distance between the focal point of the spindle body and the MTOC; this distance was markedly higher upon ectopic addition of cytATL, on average 3 times the distance measured in control embryos (Fig. 4F, 0.4 ± 0.2 µm versus 1.2 ± 0.5 µm). Overall, our spatially resolved perturbation of ER membranes reveals that the ER surrounding the spindle poles plays multiple roles in spindle architecture, including spindle size and spindle pole attachment.

**Figure 4:**
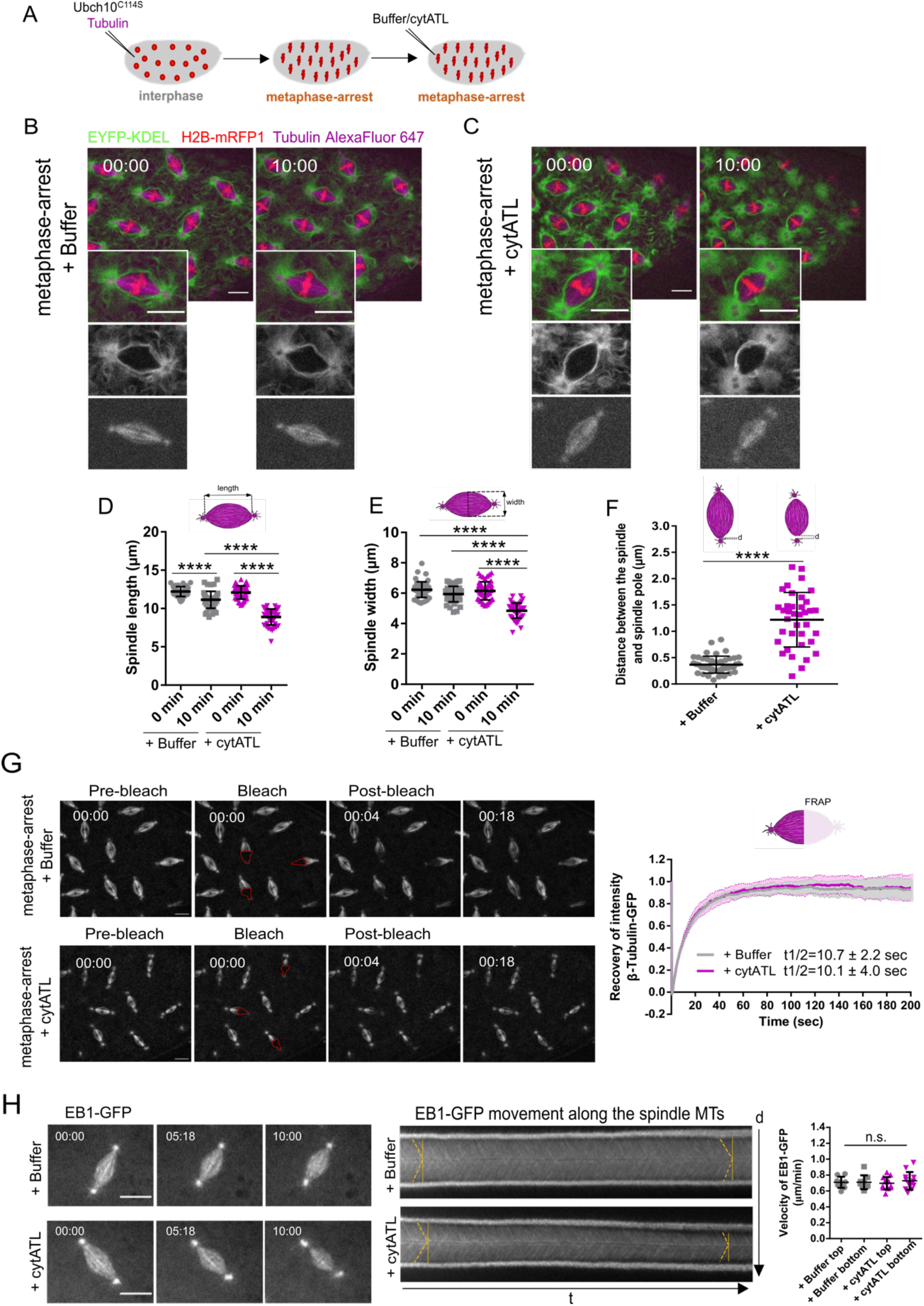
Spindle shape and function are affected upon cytATL-mediated ER disruption. **(A)** Schematics of the experimental layout: embryos were microinjected with UbcH10^C114S^ and Alexa Fluor 647-labelled Tubulin (to visualize spindle microtubules), followed by subsequent microinjection with buffer/cytATL. **(B-C)** Representative control (B) and cytATL-injected embryos (C), showing the ER (EYFP–KDEL, green), chromosomes (H2B–mRFP, red) and the spindle (magenta) at the first (t=00:00 min) and last (t=10:00 min) time points after microinjection with buffer/cytATL. Grey panels depict ER (top) and spindle (bottom) alone. Scale bar is 10 µm. **(D-E)** Quantification of spindle length (D) and width (E) in control (+Buffer, grey) and cytATL (+cytATL, magenta) embryos at the first (t=0 min) and last (t=10 min) time points of the time lapse. Asterisks depict the statistical significance derived from One-way ANOVA, multiple comparisons, with the p value adjusted to multiple comparisons (Tukey). **** p<0.0001 (N=10 embryos, n=5 nuclei). **(F)** Distance d between the focal point of the spindle and the spindle pole. Arrowheads highlight spindle pole detachment observed upon microinjection with cytATL. Asterisks depict the statistical significance derived from unpaired t-test, **** p<0.0001 (N=10 embryos, n=5 nuclei). **(G)** FRAP assay of β-tubulin–GFP in UbcH10^C114S^-arrested embryos (metaphase arrest) in control (Buffer injection) and ER disruption (+ cytATL) conditions. Right: graph depicts recovery of β-tubulin-GFP intensity after bleaching in both conditions and calculated half times of recovery (mean ± SD, N=5 embryos, n=3 nuclei). **(H)** Analysis of MT growth rate with and without ER disruption (buffer/cytATL injection) in metaphase arrested embryos. Microtubule plus-ends were monitored using a GFP-tagged EB1 transgene. Time (min:sec) is relative to microinjection with buffer/cytATL. Scale bar is 10 µm. Middle: Kymographs of EB1–GFP intensity were generated and the angles of EB1 tracks (yellow lines) were used to estimate the velocities of MT growth, shown on the right. Statistical analysis was performed using unpaired t-test, n.s. = non-significant, p>0.05 (N=4 embryos, n=3 nuclei).

Given the observed changes in spindle architecture upon impairment of spindle pole-proximal ER membranes, we next investigated whether spindle dynamics would also be affected. We used FRAP to bleach one half of the spindle in metaphase-arrested embryos and analyzed spindle microtubule turnover. We used embryos expressing β-tubulin–GFP and monitored the recovery of the fluorescence intensity after photobleaching (Fig. 4G, Suppl. Video 7). In control (buffer-injected) conditions, this analysis revealed that spindle microtubules recover very fast, with a half-time of recovery of 10.7 ± 2.2 s (Fig. 4G, grey). We next repeated the same analysis after cytATL-mediated ER disruption. In this condition, we found that MT turnover remained unaltered relative to controls (Fig. 4G, magenta). These results imply that despite the marked difference in spindle size, the dynamic behavior of MTs remains unaltered. To confirm this notion, we measured MT growth using the plus-end protein EB1 and time-lapse imaged embryos expressing a GFP-tagged EB1 protein (EB1–GFP) in control or cytATL injected embryos (Fig. 4H, left, Suppl. Video 8). Since EB1–GFP localizes to growing microtubule ends it displayed moving speckles within the spindle in time-lapse images. To estimate MT growth speed, we generated space-time projections (kymographs) along the spindle axis, which transform the continuously moving speckles to linear signals (Fig. 4H, middle). Analysis of growth speed revealed no significant differences between control and cytATL injected embryos (Fig. 4H, right). Thus, we conclude that the observed changes in spindle architecture upon ectopic addition of cytATL are not accompanied by changes of microtubule dynamics. From this insight, we hypothesized that the ER confers constraining mechanical force on the mitotic spindle, which could play a role in spindle function.

### cytATL-mediated disruption of the ER impairs mitotic spindle pulling forces

We next asked how the smaller spindles that result from acute and local perturbation of spindle pole-proximal ER membranes behave at the functional level. For this, we took advantage of a molecular tool that enables the artificial separation of sister chromatids. With this approach, fast Cohesin inactivation is achieved by the Tobacco Etch Virus (TEV) protease in embryos surviving on a modified version of Rad21 that contains TEV cleavage sites (Pauli et al. 2008). Upon microinjection of TEV protease, sister chromatid separation is observed within 1–2 minutes (Oliveira et al. 2010; Carmo et al. 2019). This tool allowed us to investigate changes in pulling force on chromatids, as poleward movement of separated sister chromatids relies on how efficiently they are pulled apart by spindle microtubules. Embryos were first arrested in metaphase, followed by ectopic addition of cytATL or buffer. 10 minutes after cytATL/buffer injection, embryos were injected with TEV protease to trigger artificial sister chromatid separation (Fig. 5A, Suppl. Video 9). Under both conditions sister chromatids separated (Fig. 5B). Analysis of chromosome movement was performed based on kymographs that display chromosome position over time (Fig. 5C). In control embryos, we found that sister chromatid separation, defined by the bifurcation in the kymograph (Fig. 5C, arrow), is elicited within ∼2 minutes upon microinjection with TEV protease while it is delayed upon ectopic addition of cytATL (Fig. 5D). Moreover, the velocity of initial poleward-movement of sister chromatids along the spindle axis, as estimated by the angle of signal bifurcation in the kymograph, is reduced upon cytATL-driven ER disruption compared to the control condition (Fig. 5E). The range of chromatid separation along the spindle axis is also reduced upon cytATL addition (Fig. 5F), which we attribute to lower separation velocity and overall shorter spindle size. After the initial separation, isolated sisters from both experimental conditions (buffer/cytATL) were equally able to engage into oscillatory movements driven by cycles of chromosome capture/detachment (Sup Fig. 3). We conclude that the short spindles imposed by acute ER disruption are still able to pull and capture chromatids. However, poleward chromatid movement occurs at slower velocity, implying a decrease in pulling forces on chromatids.

**Figure 5:**
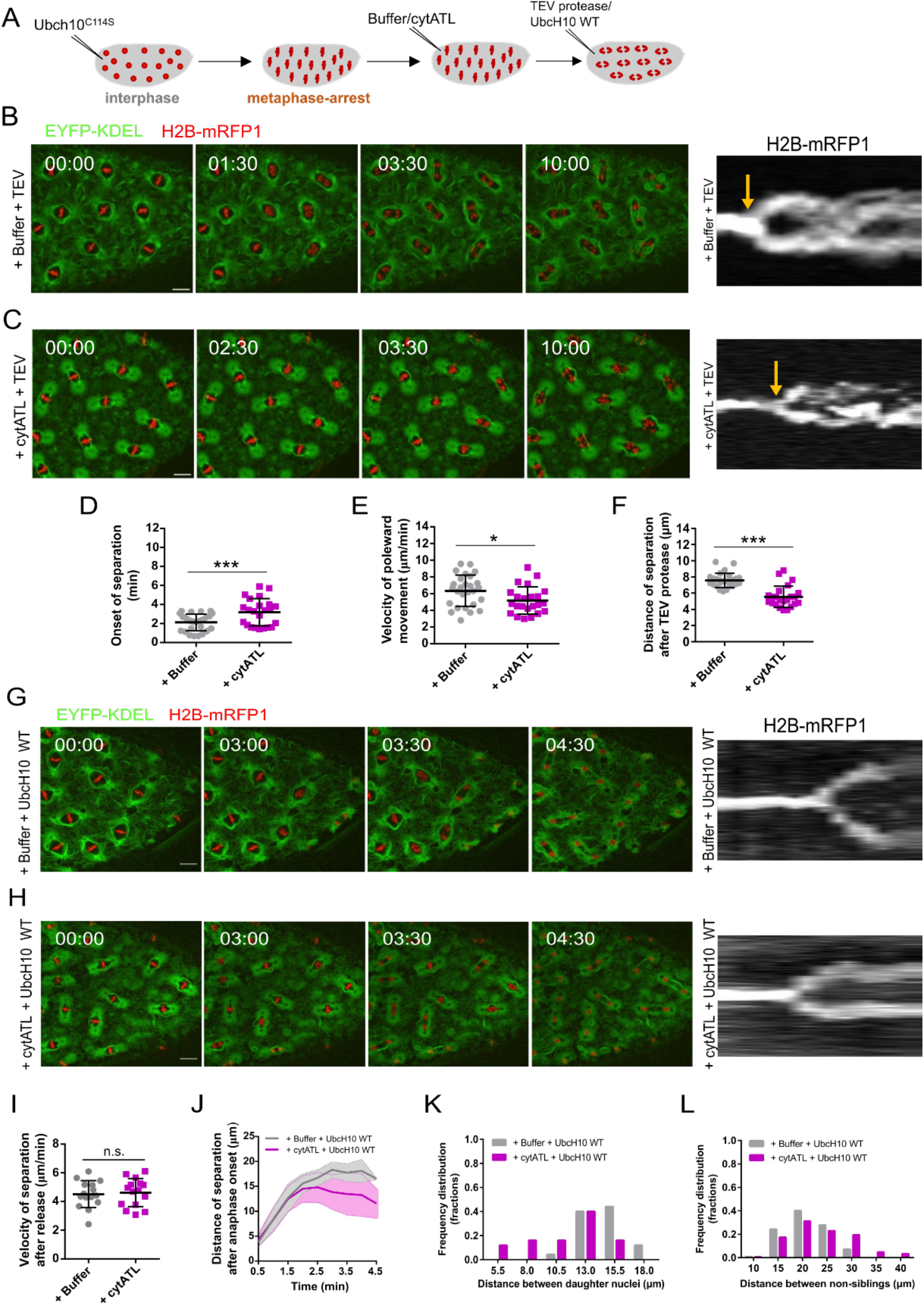
Spindle function is impaired upon cytATL-mediated ER disruption. **(A)** Experimental layout: embryos were arrested in metaphase (UbcH10^C114S^ microinjection), followed by microinjection with buffer/cytATL. After 10 min, embryos were subjected to a third microinjection with either TEV protease, to induce acute sister chromatid separation, or UbcH10^WT^ to trigger mitotic exit. Embryos surviving solely on a TEV-cleavable version of Rad21 (Cohesin) were used for these experiments. **(B-C)** Stills from embryos after microinjection with buffer/cytATL were subsequently microinjected with TEV protease to trigger cohesin cleavage (+Buffer+TEV, B; +cytATL+TEV, C); ER is labelled in green (EYFP–KDEL) and chromosomes in red (H2B–mRFP1); scale bar is 10 µm; time (min:sec) is relative to microinjection with TEV; kymographs (right panel) depict chromosome positioning over time; arrows highlight the onset of chromatid separation. **(D-F)** Quantification of the onset of separation (D), velocity of poleward movement (E) and distance of chromatids separation (F) induced by cohesin cleavage (+TEV), in conditions with intact ER (+Buffer) or disrupted ER (+cytATL). Sample size: N=5 embryos, n=5 different nuclei for each experimental condition. **(G-H)** Time lapse images of induced anaphase in unperturbed ER (G, +Buffer+UbcH10^WT^) and perturbed ER (H, +cytATL+UbcH10^WT^) conditions; ER is labelled in green (EYFP–KDEL) and chromosomes in red (H2B–mRFP1); scale bar is 10 µm; time (min:sec) is relative to microinjection with UbcH10^WT^; kymographs (right panel) depict chromosome positioning over time; **(I-J)** Quantification of the velocity of chromosome separation (H) and distance of chromosome separation (J) triggered upon artificial induction of anaphase (UbcH10^WT^) in embryos previously injected with buffer or cytATL. Sample size: N=5 embryos, n=3 different nuclei for each experimental condition. **(K-L)** Frequency distributions of distance between daughter nuclei (K) and non-sibling nuclei (L) measured 3 minutes after anaphase onset; sample size: 3-4 measurements, N=5 embryos (K) and 6 measurements, N=5 embryos (L). Asterisks refer to statistical significance, derived from unpaired t tests: * = p<0.05, ** = p<0.01, *** = p<0.001 **** = p<0.0001, n.s. = non-significant, p>0.05.

### cytATL-driven disruption of the ER affects nuclear spacing upon release from the metaphase arrest

Our previous work has revealed that the spindle pole MTOC plays a crucial role in daughter nuclei separation and nuclear spacing in the syncytial embryo (Telley et al. 2012; de-Carvalho et al. 2022). Having observed spindle pole detachment upon cyATL injection (Fig. 3), we next aimed to probe nuclear separation after disruption of the ER in embryos undergoing the changes characteristic of a normal mitotic exit. For this we took advantage of an inducible release of the metaphase-arrest reported previously (Piskadlo, et al. 2017): we induced a metaphase-arrest in division cycle 10 using the dominant-negative UbcH10^C114S^ for 5 min, followed by injection of buffer/cytATL and waited for 10 min to allow for ER disruption. Embryos were subsequently injected with a wild-type version of UbcH10 protein, which induces anaphase onset and, thus, mitotic exit in 4–8 min (Fig. 5G). Using this approach coupled to time-lapse imaging, we then generated kymographs of the chromatin signal and tracked chromosome segregation and daughter nuclei separation. This analysis revealed that sister chromatids are separated at a similar velocity during anaphase in both control and cytATL conditions (Fig. 5G-H, Suppl. Video 10). This contrasts with what we observed in embryos arrested in metaphase (+TEV cleavage of cohesin) suggesting that anaphase-specific changes may compensate for the reduced pulling forces observed upon ER disruption in metaphase. However, despite normal segregation speeds, daughter nuclei were not efficiently separated during telophase and early interphase upon ER disruption (Fig. 5H). This process is driven by the centrosome-nucleated microtubule aster (Telley et al. 2012) and, therefore, the reduced separation upon ER disruption is likely caused by the defects observed on spindle pole attachment. Furthermore, the inefficient separation led to a lower nuclear ordering in the subsequent interphase. In our control experiments, UbcH10-arrest/release approach is able to reproduce the non-sibling internuclear distance recently reported for this division cycle (cycle 11) (de-Carvalho et al. 2022). In contrast, upon ER disruption, sibling nuclear distance was shorter and the distances between non-sibling nuclei were longer when compared to the control condition (Fig. 5J-L).

In summary, here we show that ER membranes located in the vicinity of the spindle poles are critical for maintenance of proper mitotic spindle shape and function. This novel role for ER membranes is required throughout metaphase, even after unperturbed spindle assembly. Our findings highlight that the role of the ER in spindle architecture goes well beyond the phase of spindle assembly at early mitotic stages, as previously reported (Schweizer et al. 2015; Liu and Zheng 2009). We favor that this constant requirement may underlie the observed continuous remodeling of ER membranes, as evidenced by the highly dynamic behaviors of mitotic ER membranes. Cdk1 consensus sequences were found in *Drosophila* ER-shaping proteins such as Rtnl1, Spastin and Atlastin (Bergman et al. 2015), suggesting that their activity is spatiotemporally regulated throughout the cell cycle. Moreover, the human ortholog of Atlastin-1 interacts with Spastin, a microtubule-severing ATPase, within tubular ER membranes in neurons (Park et al. 2010), molecularly linking ER shaping and microtubule dynamics. However, it remains to be determined how the centrosome-proximal ER membranes maintain spindle shape in dividing tissues. Previous work suggested that membranous structures surrounding the spindle can affect molecular crowding and thereby impact on spindle assembly (Schweizer et al. 2015). Although we cannot exclude whether changes in the molecular composition are also imposed in our experiments, the finding that spindle dynamics remains largely unaltered strongly suggests this is not the case. Instead, we favor that ER membranes at the spindle poles may aid on balance of forces within spindle. Mechanical force transmission and balancing may be related to the attachment of the spindle pole MTOC and the anchoring of the microtubule aster to the nuclear envelope. This is particularly relevant during syncytial divisions, as ordered nuclear positioning needs to be established in the absence of proximal cell membrane. This spatial distribution of dividing nuclei is a crucial event for later developmental steps, such as cell size determination and gene expression patterning. Future work should address the role of the ER in other cells where nuclear positioning is critical, with direct implications on further organism development.

## Acknowledgements

We thank members of the Telley and Oliveira labs for fruitful discussions. We thank the staff of the Fly Facility, the Advanced Imaging Facility (AIF) and the Technical Support Service at the Instituto Gulbenkian de Ciência (IGC).

## Author contribution

M.A., I.A.T. and R.A.O. conceived the study; M.A. established the fly strains, designed and carried out all the experiments; A.T. and D.V.V were involved in protein production and purification; M.A., I.A.T. and R.A.O. analyzed the data; M.A. I.A.T. and R.A.O. wrote the draft manuscript; all authors edited the final manuscript; I.A.T. and R.A.O. supervised the project.

## Materials and Methods

### Fly stocks

Fly strains expressing His2Av–mRFP (Schuh et al. 2007), EYFP-ER (LaJeunesse et al. 2004), Rtnl1–GFP (Morin et al. 2001) fluorescent markers, and for induction of artificial sister chromatid separation (Pauli et al. 2008; Oliveira et al. 2010) have been previously described. A list with all stocks can be found in table 1.

**Table 1.**
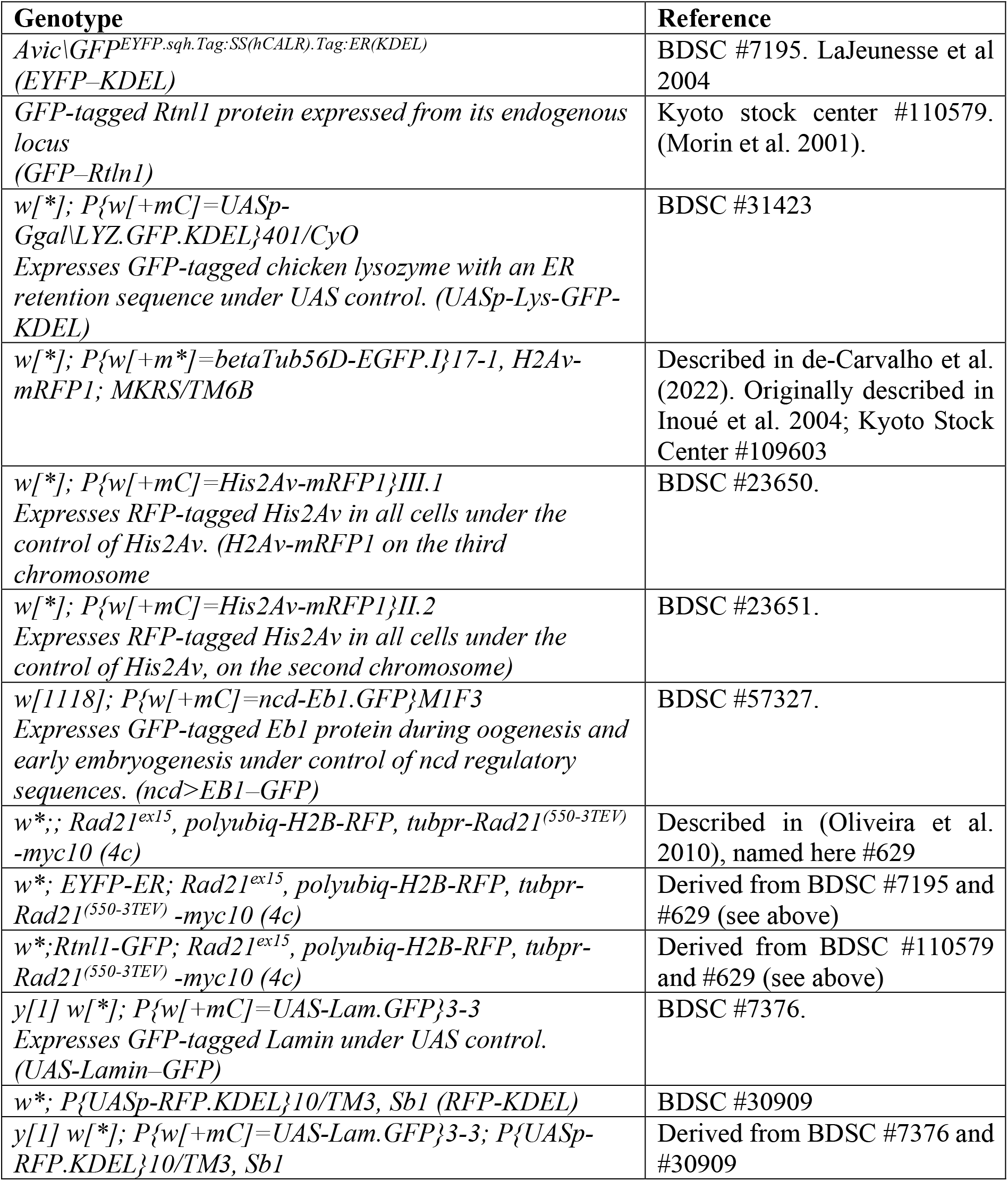
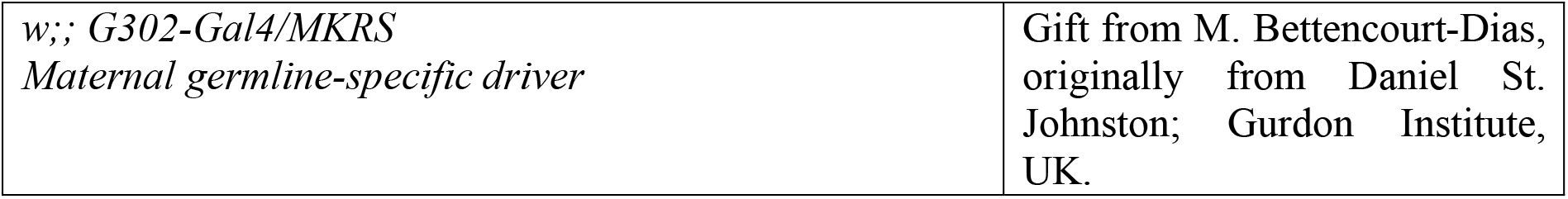
List of *Drosophila* stocks used in this study

| Genotype | Reference |
| --- | --- |
| *Avic\GFP<sup>EYFP.sqh.Tag:SS(hCALR).Tag:ER(KDEL)</sup>*<br>(*EYFP-KDEL*) | BDSC #7195. LaJeunesse et al 2004 |
| *GFP-tagged Rtnl1 protein expressed from its endogenous locus*<br>(*GFP-Rtnl1*) | Kyoto stock center #110579. (Morin et al. 2001). |
| *w[\*]; P{w[+mC]=UASp-Ggal\LYZ.GFP.KDEL}401/CyO*<br>*Expresses GFP-tagged chicken lysozyme with an ER retention sequence under UAS control. (UASp-Lys-GFP-KDEL)* | BDSC #31423 |
| *w[\*]; P{w[+m\*]=betaTub56D-EGFP.I}17-1, H2Av-mRFP1; MKRS/TM6B* | Described in de-Carvalho et al. (2022). Originally described in Inoué et al. 2004; Kyoto Stock Center #109603 |
| *w[\*]; P{w[+mC]=His2Av-mRFP1}III.1*<br>*Expresses RFP-tagged His2Av in all cells under the control of His2Av. (H2Av-mRFP1 on the third chromosome)* | BDSC #23650. |
| *w[\*]; P{w[+mC]=His2Av-mRFP1}II.2*<br>*Expresses RFP-tagged His2Av in all cells under the control of His2Av, on the second chromosome)* | BDSC #23651. |
| *w[1118]; P{w[+mC]=ncd-Eb1.GFP}M1F3*<br>*Expresses GFP-tagged Eb1 protein during oogenesis and early embryogenesis under control of ncd regulatory sequences. (ncd>EB1-GFP)* | BDSC #57327. |
| *w\*;; Rad21<sup>ex15</sup>, polyubiq-H2B-RFP, tubpr-Rad21<sup>(550-3TEV)</sup>-myc10 (4c)* | Described in (Oliveira et al. 2010), named here #629 |
| *w\*; EYFP-ER; Rad21<sup>ex15</sup>, polyubiq-H2B-RFP, tubpr-Rad21<sup>(550-3TEV)</sup>-myc10 (4c)* | Derived from BDSC #7195 and #629 (see above) |
| *w\*; Rtnl1-GFP; Rad21<sup>ex15</sup>, polyubiq-H2B-RFP, tubpr-Rad21<sup>(550-3TEV)</sup>-myc10 (4c)* | Derived from BDSC #110579 and #629 (see above) |
| *y[1] w[\*]; P{w[+mC]=UAS-Lam.GFP}3-3*<br>*Expresses GFP-tagged Lamin under UAS control. (UAS-Lamin-GFP)* | BDSC #7376. |
| *w\*; P{UASp-RFP.KDEL}10/TM3, Sb1 (RFP-KDEL)* | BDSC #30909 |
| *y[1] w[\*]; P{w[+mC]=UAS-Lam.GFP}3-3; P{UASp-RFP.KDEL}10/TM3, Sb1* | Derived from BDSC #7376 and #30909 |

### Microinjections

Microinjections were performed as previously described in (Carmo et al. 2019), using an Eppendorf FemtoJet Microinjector controller and commercially available needles (Eppendorf™ Femtotips™ Diameter (Metric) Inner: 0.5 μm, 11883991). Briefly, dechorionated embryos (1–1.5 h old) were aligned and glued onto a #1.5 coverslip (24×40 mm), dried at room temperature for 13 min, and subsequently covered in halocarbon oil. Buffer/protein/chemicals were microinjected at the following concentrations: Buffer (10 mM HEPES, 100 mM KCl, 1 mM MgCl_2_, 10% glycerol), UbcH10^C114S^ (30-40 mg/ml), p27 (4 mg/ml), TEV protease (18 mg/ml), cytATL (60 mg/ml), UbcH10 wildtype (60 mg/ml). Porcine Tubulin labelled with AlexaFluor 647 (Tubulin HILyte Fluor™ 647 labelled, cat. # 1L670M, 20 µg; Cytoskeleton) was reconstituted in freshly prepared and filtered 1X BRB80 buffer (80 mM PIPES, 1 mM MgCl_2_, 1 mM EGTA, pH 7.8) to a final concentration of 2.5 mg/ml, immediately flash frozen in liquid nitrogen and stored at - 80°C. To visualize spindle microtubules upon metaphase-arrest, labelled Tubulin was mixed with UbcH10^C114S^ protein at 1:1 ratio (at a final concentration of 1.25 mg/ml). To visualize spindle microtubules during nuclear division cycles, Tubulin was used at 2.5 mg/ml.

### Protein purification

UbcH10^C114S^, p27, TEV protease were purified as previously described (Piskadlo et al. 2017; Oliveira et al. 2010). For cytATL, the bacterial expression plasmid containing *D. melanogaster* cytATL sequence (pET28a-His_6_-cytATL; kindly provided by Junjie Hu, Nankai University, China) was used to transform *E. coli* BL21 cells. Bacterial cells from an overnight-grown starting culture were grown in 1L LB medium, supplemented with Kanamycin (50 µg/ml), and incubated at 37°C. Protein expression was induced by adding 0.2 mM IPTG once the O.D._600_ was at 0.8. The culture was grown at 18°C overnight and bacteria were harvested by centrifugation at 4200 rpm for 45 min, at 4°C. For protein purification, the following buffers were prepared freshly and filtered: 0.1 M KPi at pH 7.2; wash buffer (50 mM KPi pH 7.2, 400 mM NaCl, 1 mM ß-Mercaptoethanol, 7 mM imidazole), lysis buffer (50 mM KPi pH7.2, 400 mM NaCl, 1 mM ß-Mercaptoethanol, 7 mM imidazole, 0.1% Triton X-100 supplemented with 1 tablet of protease inhibitors (Pierce™ Protease Inhibitor Mini Tablets, EDTA-free, reference A32955, Thermo Scientific) and DNAseI), elution buffer 1 (50 mM KPi pH7.2, 400 mM NaCl, 1 mM ß-Mercaptoethanol, 400 mM imidazole, elution buffer 2 (50 mM KPi pH7.2, 400 mM NaCl, 1 mM ß-Mercaptoethanol, without imidazole). Bacterial pellets were re-suspended in 10 ml lysis buffer, adding DNase and a protease inhibitor tablet onto the pellet directly. The lysate was then passed through a French press system (Emulsiflex C5 High Pressure Homogeneizer). Protein extract was spun at 16000 rpm for 45 min at 4°C, and the pellet was discarded. Ni sepharose beads (2-3 ml; GE Healthcare) were added into a 50 ml falcon tube. Protein extract was incubated with beads for 1h at 4°C with gentle agitation. Beads were washed 3 times with wash buffer (each round at 700 g for 5 min) and packed into a COLUMN PD-10 EMPTY (GE Healthcare). The protein was eluted in distinct fractions with increasing concentrations of imidazole (80, 150, 300, 400 mM). The two fractions eluted with 300 and 400 mM imidazole were pooled together and dialyzed overnight at 4°C using a cassette (Slide-A-Lyzer Dialysis Cassettes 7 KDa MWCO, 12 ml, Life Technologies) into a 1 L volume of cytATL storage buffer (10 mM HEPES, 100 mM KCl, 1 mM MgCl_2_, 10% glycerol). To concentrate the purified cytATL protein we used Amicon ultra centrifugal filter 15 mL with a 10 kDa cut-off (Millipore) and protein concentration was quantified in a nanodrop using the A280.

### Microscopy

Time-lapse movies of live embryos were obtained using Confocal Z-series stacks with a Yokogawa CSU-X Spinning Disk confocal, mounted on a Leica DMi8 microscope, with a 63x 1.3NA glycerine immersion objective, using the 488 nm and 561 nm laser lines and a Andor iXon Ultra EMCCD 1024×1024 camera. The system was controlled with Metamorph software (Molecular Devices). For the sequential microinjection experiments, we used 30 sec time points and a total of 10 min for each time-lapse acquisition, with 0.4-0.5 µm z-step size and 15 slices

### Quantitative imaging analysis

To quantify the area and perimeter of the ER exclusion zone and of the spindle, a single z slice corresponding to the middle plane of each nucleus was used. All the measurements were performed using the segmented line tool in Fiji (yellow shapes in Fig. 1A, insets), for each channel (ER, spindle) separately.

### FRAP assays

FRAP experiments were performed using Andor’s Mosaic system with a 470 nm laser, using 0.25 s (Lys–GFP–KDEL expressing embryos) or 0.5 sec time points (Rtnl1-GFP expressing embryos) and a single z plane was acquired. Collective movement of spindles in the embryo was corrected using the *stackreg* plugin in Fiji (Thévenaz et al. 1998). The first time point was used as a prebleach reference of fluorescence intensity and bleaching was set to the second time point. Raw fluorescence intensity values were extracted from the time-lapse movies with the plot z-axis profile option in Fiji. FRAP recovery curves were analyzed using the easyFRAPweb tool. Briefly, three different regions of interest (ROIs) were measured over time: the bleached region (circle with 42 pixels diameter), the entire ER compartment of a single nucleus (variable size), and the background (region outside the embryo, circle with 95 pixels diameter). ROI positioning was kept constant throughout the time series (this led to occasional inflow of Lys–GFP–KDEL-labeled vesicles, which may slightly overestimate the mobile pool in this strain).

### ER fluorescence intensity

ER mean intensity of control-(+Buffer) and cytATL-injected embryos was measured for the last time point (t=10 min) of the time-lapse. A circle ROI with the same area was used to measure the mean intensity at the spindle pole or the equator regions in the ER channel. Relative signal dispersion was calculated by the ratio between the standard deviation (stdev) and the mean fluorescence intensity (mean).

### Quantification of MT growth and sister chromatid movement

Analysis of EB1 dynamics and chromatid poleward movement was performed based on kymographs, created using the FIJI kymograph plug-in (written by J. Rietdorf and A. Seitz, EMBL, Heidelberg, Germany). We estimated speed by measuring the angle of the linear signals (EB1–GFP for MT growth, histone H2B–mRFP1 for chromatid movement) in the kymographs. Quantification of chromosome movement (Supp. Fig. 3A) was performed as previously described (Mirkovic et al. 2015). Briefly, H2B–mRFP1 was imaged at 30 sec intervals and images were segmented to select the chromosomal regions, based on an automatic threshold (set in the last frame, 10 min after TEV injection). For each binary-images movie, a walking average of 3 frames was produced (using kymograph plug-in, written by J. Rietdorf and A. Seitz, EMBL, Heidelberg, Germany) creating a merged image in which the intensity is proportional to the overlap between consecutive frames. Intensity profiles were used to estimate the percentage of non-overlapping, 2-frame overlap and 3-frame overlap pixels. The area occupied by sister chromatids (Supp. Fig. 3B) was calculated using a macro that filters, creates a mask, and subsequently fits a convex hull algorithm enclosing all the sister chromatids. Then, a spline connecting all the sister chromatids is created. The area of this fitted spline is measured at each time point, estimating the area occupied by sister chromatids at each time frame.

### Distance between sibling and non-sibling nuclei

Measurements of distance between sibling (daughter) and non-sibling nuclei were done using the line tool in Fiji, 3 min after anaphase onset (counting from the time point of sister chromatid separation). On average, five pairs of daughter nuclei and six non-sibling (neighboring) nuclei were measured. Graphic representation was performed using Prism seven software (RRID:SCR_002798, GraphPad, La Jolla, CA).

## Supplemental Material

**Supplementary Figure 1:**
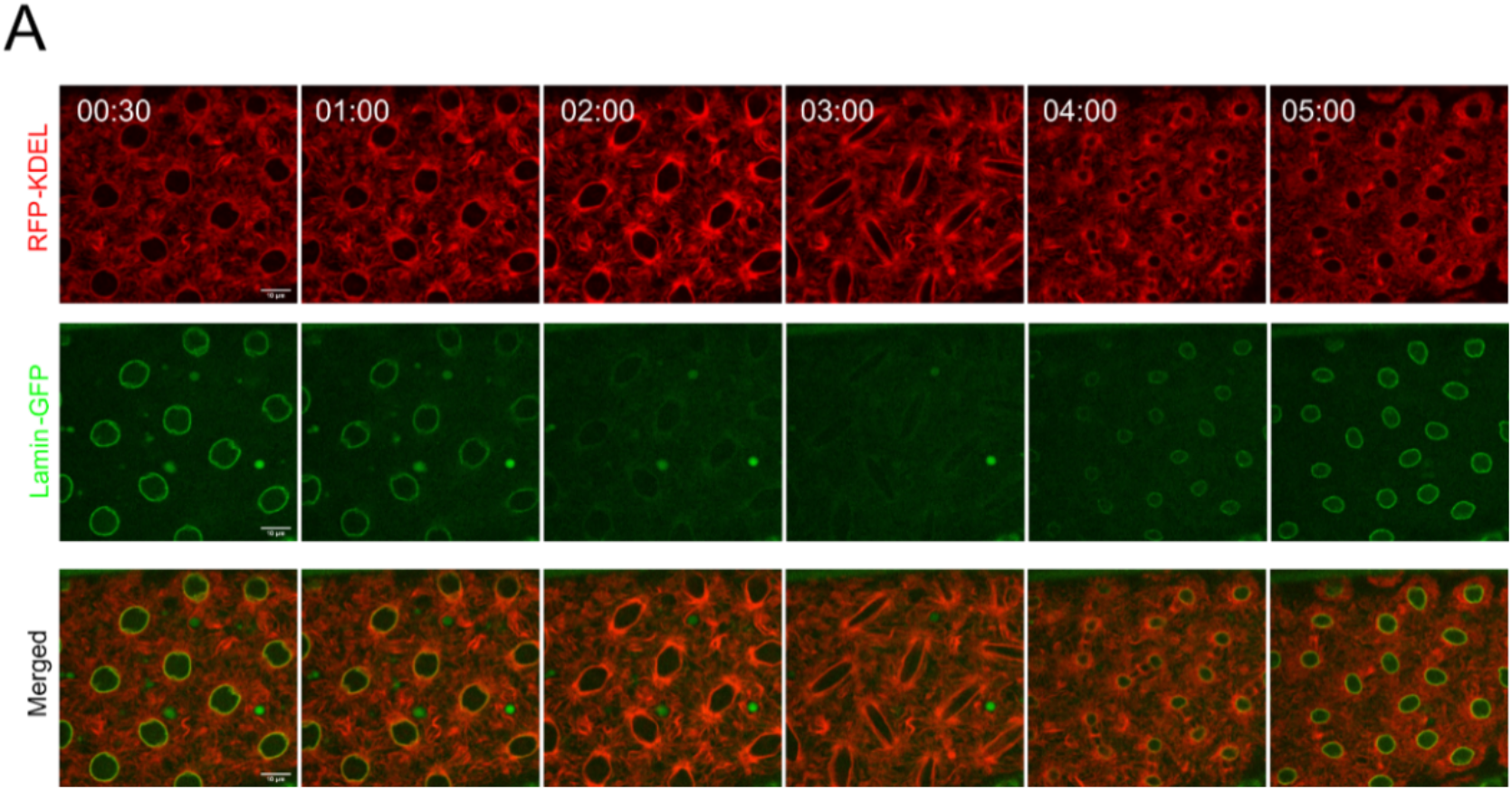
Time-lapse images of nuclear division cycle in embryos expressing RFP–KDEL (ER, red) and Lamin B–GFP (nuclear envelope, green) under the control of the G302-Gal4 maternal driver. The ER extends from the nuclear envelope as indicated by the merged image (t=00:30, merged, yellow). As nuclei enter mitosis (t=01:00 to 03:00, green), a partially disassembled Lamin B envelope can be observed while the ER is present continuously throughout the cell cycle (red). The nuclear envelope is reassembled at telophase (t=04:00, green).

**Supplementary Figure 2:**
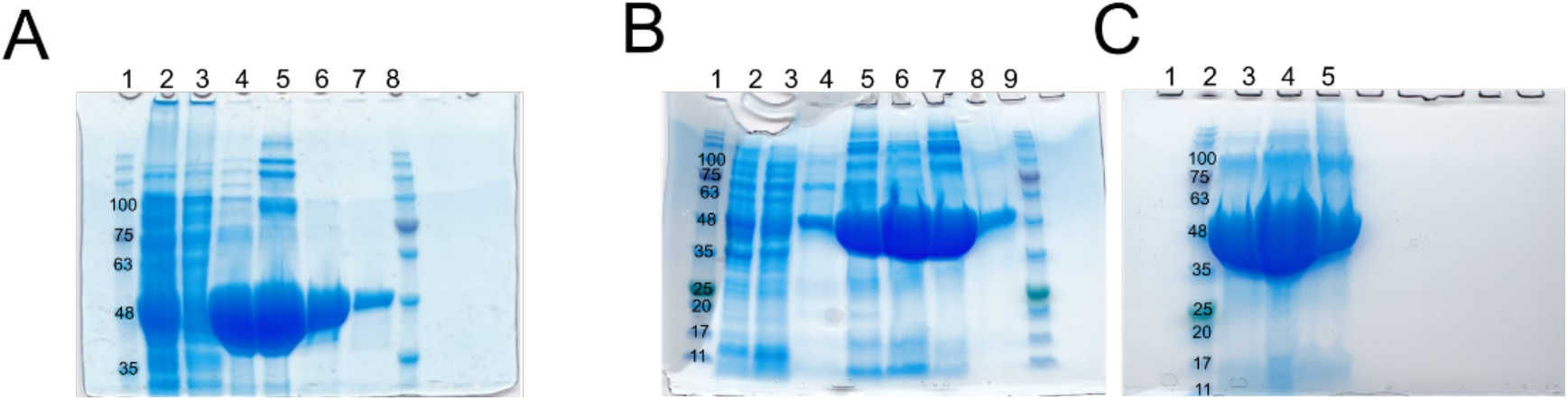
Expression and purification of *D. melanogaster* cytoplasmic domain of Atlastin (His_6_-cytATL). **(A)** First His6-cytATL protein purification batch using Ni sepharose beads (see Materials and Methods for detailed protocol). 1 – Marker (NZY tech protein marker III). 2 – Supernatant after lysis. 3 – Supernatant after incubation with beads. 4, 5, 6 and 7 are fraction eluted with 80, 150, 300 and 400 mM imidazole respectively. Pooled fractions (6 and 7) were dialyzed overnight into storage buffer, concentrated to 60 mg/ml, flash frozen in liquid nitrogen. Theoretical His_6_-cytATL protein size is 48 kDa. **(B)** Second purification batch using the same protocol as in (A), but with increased volume of beads up to 3 ml. Rows: 1 – Marker, 2 – supernatant after lysis, 3 – supernatant after incubation with beads. 4, 5, 6, 7, 8 – fraction eluted with 40, 80, 150, 300 and 400 mM imidazole, respectively. 9 – Marker. **(C)** Fractions 6 (lane 3 and 4 with 5 and 10 µl purified protein, respectively) and 8 (lane 5 with 5 µl purified protein) shown in B.

**Supplementary Figure 3:**
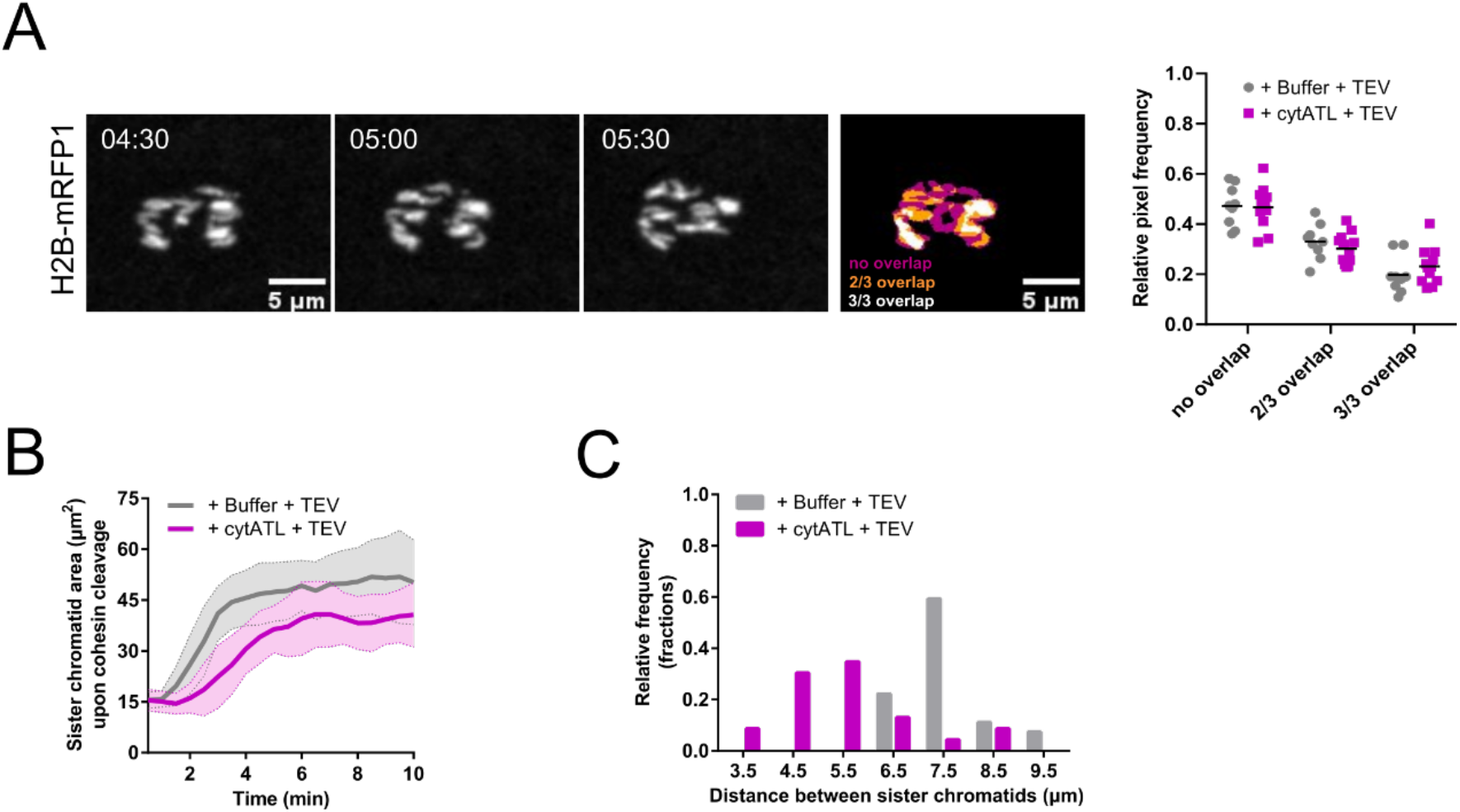
Quantification of oscillatory movements upon sister chromatid separation induced by TEV-mediated cleavage of Cohesin. **(A)** Quantification of the oscillatory behavior of separated sister chromatids (related to Fig. 5, upper panel). The movement of sister chromatids was estimated by measuring the signal overlap between three consecutive frames (no overlap – magenta, 2 out of 3 frames overlap – orange, 3 out of 3 frames overlap – white). Relative pixel frequency for each category is quantified, displaying no significant difference between the two conditions (Buffer – grey, cytATL – magenta). Statistical analysis using N=5 embryos, n=5 nuclei per embryo, using unpaired t-test, two-sided. revealed no statistical differences between buffer and cytATL injected embryos (p>0.05) **(B)** Quantification of the area occupied by sister chromatid upon TEV-cleavage of Cohesin, depicting a delay in the onset of sister chromatid separation and a lower area occupied in space for cytATL (magenta) compared to control embryos (grey). **(C)** Frequency distribution of distance between sister chromatids indicating an increased frequency of measurements ≤ 5.5 µm for cytATL embryos (magenta).

**Suppl. Video 1**: Syncytial nuclear divisions in *Drosophila* embryos. ER is labeled with EYFP–KDEL (left, green) or GFP–Rtnl1 (right, green) together with Histone H2B– mRFP1 (red) after microinjection with Tubulin AlexaFluor-647 (magenta, to label microtubules). Times are relative to injection. Scale bar is 10 µm. Acquisition frame rate is 2 min^-1^. Display frame rate is 3 s^-1^.

**Suppl. Video 2**: FRAP experiment in embryos containing GFP–Rtnl1 (black), after microinjection with 4 mg/ml of Cdk1 inhibitor p27 to induce an interphase arrest. Scale bar is 10 μm. Time in seconds is relative to pre-bleaching. Bleaching was performed at the second time point, followed by monitoring of recovery of fluorescence intensity over time. Acquisition frame rate is 2 s^-1^. Display frame rate is 3 s^-1^.

**Suppl. Video 3**: FRAP experiment on the ER in embryos containing GFP–Rtnl1. Microinjection with 40 mg/ml of a dominant-negative form of the human E2 ubiquitin-conjugating enzyme (UbcH10^C114S^) was performed to induce a metaphase arrest. Scale bar is 10 μm. Time in seconds is relative to pre-bleaching. Bleaching was performed at the second time point, followed by monitoring of recovery of fluorescence intensity over time. Acquisition frame rate was 2 s^-1^. Frame rate is 3 frames per second.

**Suppl. Video 4**: FRAP experiment on the ER in embryos containing Lys–GFP–KDEL. Microinjection with 40 mg/ml of a dominant-negative form of the human E2 ubiquitin-conjugating enzyme (UbcH10^C114S^) was performed to induce a metaphase arrest. Scale bar is 10 μm. Time in seconds is relative to pre-bleaching. Bleaching was performed at the second time point, followed by monitoring of recovery of fluorescence intensity over time. Acquisition frame rate was 4 s^-1^. Frame rate is 3 frames per second.

**Suppl. Video 5:** Embryos containing EYFP–KDEL, after microinjection with 40 mg/ml of UbcH10^C114S^ to induce a metaphase arrest, and subsequently microinjected with buffer (left) or cytATL (right, 60 mg/ml). Chromosomes are labeled by H2B–mRFP1 (red). Scale bar is 10 μm. Time (min:s) is relative to the time of UbcH10^C114S^ microinjection. Acquisition frame rate was 2 min^-1^. Display frame rate is 3 s^-1^.

**Suppl. Video 6**: Embryos containing EYFP–KDEL, after microinjection with 40 mg/ml of UbcH10^C114S^, together with Tubulin AlexaFluor-647, to induce a metaphase arrest, and subsequently microinjected with buffer (left) or cytATL protein (right, 60 mg/ml). Chromosomes are labeled by H2B–mRFP1 (red). Scale bar is 10 μm. Time (min:s) is relative to the time of microinjection with buffer/cytATL. Acquisition frame rate was 2 s^-1^. Display frame rate is 3 s^-1^.

**Suppl. Video 7**: FRAP experiment on half of the spindle after microinjection with 40 mg/ml of UbcH10^C114S^, to induce a metaphase arrest, and subsequently microinjected with buffer (left) or cytATL protein (right, 60 mg/ml). Spindle microtubules are labeled with β-Tubulin–GFP. Scale bar is 10 μm. Time (min:s) is relative to pre-bleaching. Acquisition frame rate was 2 s^-1^. Display frame rate is 20 s^-1^.

**Suppl. Video 8**: Microtubule growth dynamics in embryos after microinjection with 40 mg/ml of UbcH10^C114S^, to induce a metaphase arrest, and subsequently microinjected with buffer (top) or cytATL protein (bottom, 60 mg/ml). Spindle microtubules are labeled with β-Tubulin–GFP. Scale bar is 10 μm. Time (min:s) is relative to pre-bleaching. Bleaching was performed at the second time point. Acquisition frame rate was 1 s^-1^. Display frame rate is 40 s^-1^.

**Suppl. Video 9**: Embryos containing only TEV-cleavable Cohesin, after microinjection with 40 mg/ml of UbcH10^C114S^ to induce a metaphase arrest, followed by subsequent microinjection with buffer (left) or cytATL (right). A third microinjection with TEV protease was performed to induce sister chromatid separation. ER is labeled with EYFP– KDEL (green) and chromosomes with H2B–mRFP1 (red). Scale bar is 10 μm. Time (min:s) are relative to microinjection with TEV protease. Acquisition frame rate was 2 min^-1^. Display frame rate is 3 s^-1^.

**Suppl. Video 10**: Embryo after microinjection with 40 mg/ml of UbcH10^C114S^ to induce a metaphase arrest, followed by a subsequent microinjection with buffer (left) or cytATL (right). A microinjection with the wild-type version of UbcH10 (UbcH10 WT) induced mitotic exit. ER is labeled with EYFP–KDEL (green) and chromosomes with H2B– mRFP1 (red). Scale bar is 10 μm. Time (min:s) is relative to the time of UbcH10 WT microinjection. Acquisition frame rate was 2 min^-1^. Display frame rate is 3 s^-1^.

## References

Bergman, Zane J., Justin D. McLaurin, Anthony S. Eritano, Brittany M. Johnson, Amanda Q. Sims, and Blake Riggs. 2015. “Spatial Reorganization of the Endoplasmic Reticulum during Mitosis Relies on Mitotic Kinase Cyclin a in the Early Drosophila Embryo.” PLoS ONE 10 (2): 1–23. https://doi.org/10.1371/journal.pone.0117859.

Bobinnec, Yves, Christiane Marcaillou, Xavier Morin, and Alain Debec. 2003. “Dynamics of the Endoplasmic Reticulum during Early Development of Drosophila Melanogaster.” Cell Motility and the Cytoskeleton 54 (3): 217–25. https://doi.org/10.1002/cm.10094.

Carlton, Jeremy G., Hannah Jones, and Ulrike S. Eggert. 2020. “Membrane and Organelle Dynamics during Cell Division.” Nature Reviews Molecular Cell Biology, 1–16. https://doi.org/10.1038/s41580-019-0208-1.

Carmo, Catarina, Margarida Araújo, and Raquel A. Oliveira. 2019. “Microinjection Techniques in Fly Embryos to Study the Function and Dynamics of SMC Complexes.” SMC Complexes. Part of the Methods in Molecular Biology Book Series (MIMB, Volume 2004) 2004: 1–32. https://doi.org/10.1007/978-1-4939-9520-2_19).

Champion, Lysie, Monika I. Linder, and Ulrike Kutay. 2017. “Cellular Reorganization during Mitotic Entry.” Trends in Cell Biology 27 (1): 26–41. https://doi.org/10.1016/j.tcb.2016.07.004.

Civelekoglu-Scholey, Gul, Li Tao, Ingrid Brust-Mascher, Roy Wollman, and Jonathan M. Scholey. 2010. “Prometaphase Spindle Maintenance by an Antagonistic Motor-Dependent Force Balance Made Robust by a Disassembling Lamin-B Envelope.” Journal of Cell Biology 188 (1): 49–68. https://doi.org/10.1083/jcb.200908150.

de-Carvalho, Jorge, Sham Tlili, Lars Hufnagel, Timothy E. Saunders, and Ivo A. Telley. 2022. “ Aster Repulsion Drives Short-Ranged Ordering in the Drosophila Syncytial Blastoderm.” Development 149 (2). https://doi.org/10.1242/dev.199997.

Diaz, Ulises, Zane J. Bergman, Brittany M. Johnson, Alia R. Edington, Matthew A. De Cruz, Wallace F. Marshall, and Blake Riggs. 2019. “Microtubules Are Necessary for Proper Reticulon Localization during Mitosis.” PLoS ONE 14 (12): 1–25. https://doi.org/10.1371/journal.pone.0226327.

Dumont, Sophie, and Timothy J. Mitchison. 2009. “Compression Regulates Mitotic Spindle Length by a Mechanochemical Switch at the Poles.” Current Biology 19 (13): 1086–95. https://doi.org/10.1016/j.cub.2009.05.056.

Espadas, Javier, Diana Pendin, Rebeca Bocanegra, Artur Escalada, Giulia Misticoni, Tatiana Trevisan, Ariana Velasco del Olmo, et al. 2019. “Dynamic Constriction and Fission of Endoplasmic Reticulum Membranes by Reticulon.” Nature Communications 10 (1): 1–11. https://doi.org/10.1038/s41467-019-13327-7.

Ferrandiz, Nuria, Laura Downie, Georgina P Starling, and Stephen J Royle. 2022. “Endomembranes Promote Chromosome Missegregation by Ensheathing Misaligned Chromosomes.” JCB, 2021.04.23.441091. https://doi.org/10.1101/2021.04.23.441091.

Frescas, David, Manos Mavrakis, Holger Lorenz, Robert DeLotto, and Jennifer Lippincott-Schwartz. 2006. “The Secretory Membrane System in the Drosophila Syncytial Blastoderm Embryo Exists as Functionally Compartmentalized Units around Individual Nuclei.” Journal of Cell Biology 173 (2): 219–30. https://doi.org/10.1083/jcb.200601156.

Goyal, Uma, and Craig Blackstone. 2013. “Untangling the Web: Mechanisms Underlying ER Network Formation.” Biochimica et Biophysica Acta - Molecular Cell Research 1833 (11): 2492–98. https://doi.org/10.1016/j.bbamcr.2013.04.009.

Kutay, Ulrike, Ramona Jühlen, and Wolfram Antonin. 2021. “Mitotic Disassembly and Reassembly of Nuclear Pore Complexes.” Trends in Cell Biology 31 (12): 1019–33. https://doi.org/10.1016/j.tcb.2021.06.011.

LaJeunesse, Dennis Richard, Stephanie M. Buckner, Jeffrey Lake, Charles Na, Amanda Pirt, and Kathryn Fromson. 2004. “Three New Drosophila Markers of Intracellular Membranes.” BioTechniques 36 (5): 784–90. https://doi.org/10.2144/04365st01.

Liu, Zhonghua, and Yixian Zheng. 2009. “A Requirement for Epsin in Mitotic Membrane and Spindle Organization.” Journal of Cell Biology 186 (4): 473–80. https://doi.org/10.1083/jcb.200902071.

Ma, Li, Ming-Ying Tsai, Shusheng Wang, Bingwen Lu, Rong Chen, John R Yates Iii, Xueliang Zhu, and Yixian Zheng. 2009. “Requirement for Nudel and Dynein for Assembly of the Lamin B Spindle Matrix.” Nature Cell Biology 11 (3): 247–56. https://doi.org/10.1038/ncb1832.

Maiato, Helder, Ana Gomes, Filipe Sousa, and Marin Barisic. 2017. “Mechanisms of Chromosome Congression during Mitosis.” Biology 6 (1): 13. https://doi.org/10.3390/biology6010013.

Merta, Holly, Jake W. Carrasquillo Rodríguez, Maya I. Anjur-Dietrich, Tevis Vitale, Mitchell E. Granade, Thurl E. Harris, Daniel J. Needleman, and Shirin Bahmanyar. 2021. “Cell Cycle Regulation of ER Membrane Biogenesis Protects against Chromosome Missegregation.” Developmental Cell, 1–16. https://doi.org/10.1016/j.devcel.2021.11.009.

Mierzwa, Beata, and Daniel W. Gerlich. 2014. “Cytokinetic Abscission: Molecular Mechanisms and Temporal Control.” Developmental Cell 31 (5): 525–38. https://doi.org/10.1016/j.devcel.2014.11.006.

Mirkovic, Mihailo, Lukas H. Hutter, Béla Novák, and Raquel A. Oliveira. 2015. “Premature Sister Chromatid Separation Is Poorly Detected by the Spindle Assembly Checkpoint as a Result of System-Level Feedback.” Cell Reports 13 (3): 469–78. https://doi.org/10.1016/j.celrep.2015.09.020.

Morin, Xavier, Richard Daneman, Michael Zavortink, and William Chia. 2001. “A Protein Trap Strategy to Detect GFP-Tagged Proteins Expressed from Their Endogenous Loci in Drosophila.” Proceedings of the National Academy of Sciences of the United States of America 98 (26): 15050–55. https://doi.org/10.1073/pnas.261408198.

Nourbakhsh, Kimya, Amy Ferrecccio, and Smita Yadav. 2021. “TAOK2 Is an ER Kinase That Catalyzes the Dynamic Tethering of ER to Microtubules.” The FASEB Journal 35 (S1): 1–13. https://doi.org/10.1096/fasebj.2021.35.s1.02216.

Oliveira, R A, R S Hamilton, A Pauli, I Davis, and K Nasmyth. 2010. “Cohesin Cleavage and Cdk Inhibition Trigger Formation of Daughter Nuclei.” Nat Cell Biol 12 (2): 185–92. https://doi.org/10.1038/ncb2018.

Oliveira, Raquel A., Russell S. Hamilton, Andrea Pauli, Ilan Davis, and Kim Nasmyth. 2010. “Cohesin Cleavage and Cdk Inhibition Trigger Formation of Daughter Nuclei.” Nature Cell Biology 12 (2): 185–92. https://doi.org/10.1038/ncb2018.

Park, Seong H., Peng Peng Zhu, Rell L. Parker, and Craig Blackstone. 2010. “Hereditary Spastic Paraplegia Proteins REEP1, Spastin, and Atlastin-1 Coordinate Microtubule Interactions with the Tubular ER Network.” Journal of Clinical Investigation 120 (4): 1097–1110. https://doi.org/10.1172/JCI40979.

Pauli, Andrea, Friederike Althoff, Raquel A. Oliveira, Stefan Heidmann, Oren Schuldiner, Christian F. Lehner, Barry J. Dickson, and Kim Nasmyth. 2008. “Cell-Type-Specific TEV Protease Cleavage Reveals Cohesin Functions in Drosophila Neurons.” Developmental Cell 14 (2): 239–51. https://doi.org/10.1016/j.devcel.2007.12.009.

Petry, Sabine. 2016. “Mechanisms of Mitotic Spindle Assembly.” Annual Review of Biochemistry 85 (1): 659–83. https://doi.org/10.1146/annurev-biochem-060815-014528.

Piskadlo, Ewa, Alexandra Tavares, and Raquel A. Oliveira. 2017. “Metaphase Chromosome Structure Is Dynamically Maintained by Condensin I-Directed DNA (de)Catenation.” ELife 6: 1–22. https://doi.org/10.7554/eLife.26120.

Schlaitz, Anne Lore, James Thompson, Catherine C.L. Wong, John R. Yates, and Rebecca Heald. 2013. “REEP3/4 Ensure Endoplasmic Reticulum Clearance from Metaphase Chromatin and Proper Nuclear Envelope Architecture.” Developmental Cell 26 (3): 315–23. https://doi.org/10.1016/j.devcel.2013.06.016.

Schuh, Melina, Christian F. Lehner, and Stefan Heidmann. 2007. “Incorporation of Drosophila CID/CENP-A and CENP-C into Centromeres during Early Embryonic Anaphase.” Current Biology 17 (3): 237–43. https://doi.org/10.1016/j.cub.2006.11.051.

Schweizer, Nina, Nisha Pawar, Matthias Weiss, and Helder Maiato. 2015. “An Organelle-Exclusion Envelope Assists Mitosis and Underlies Distinct Molecular Crowding in the Spindle Region.” Journal of Cell Biology 210 (5): 695–704. https://doi.org/10.1083/jcb.201506107.

Shibata, Yoko, Christiane Voss, Julia M. Rist, Junjie Hu, Tom A. Rapoport, William A. Prinz, and Gia K. Voeltz. 2008. “The Reticulon and Dp1/Yop1p Proteins Form Immobile Oligomers in the Tubular Endoplasmic Reticulum.” Journal of Biological Chemistry 283 (27): 18892–904. https://doi.org/10.1074/jbc.M800986200.

Smyth, Jeremy T., Todd A. Schoborg, Zane J. Bergman, Blake Riggs, and Nasser M. Rusan. 2015. “Proper Symmetric and Asymmetric Endoplasmic Reticulum Partitioning Requires Astral Microtubules.” Open Biology 5 (8). https://doi.org/10.1098/rsob.150067.

Snapp, Erik L, Takako Iida, David Frescas, Jennifer Lippincott-schwartz, Mary A Lilly, and Drosophila Stocks. 2004. “Connectivity in Drosophila Ovarian Cysts” 15 (October): 4512–21. https://doi.org/10.1091/mbc.E04.

Telley, Ivo A., Imre Gáspár, Anne Ephrussi, and Thomas Surrey. 2012. “Aster Migration Determines the Length Scale of Nuclear Separation in the Drosophila Syncytial Embryo.” Journal of Cell Biology 197 (7): 887–95. https://doi.org/10.1083/jcb.201204019.

Thévenaz, Philippe, Urs E. Ruttimann, and Michael Unser. 1998. “A Pyramid Approach to Subpixel Registration Based on Intensity.” IEEE Transactions on Image Processing 7 (1): 27–41. https://doi.org/10.1109/83.650848.

Tsai, Ming Ying, Shusheng Wang, Jill M. Heidinger, Dale K. Shumaker, Stephen A. Adam, Robert D. Goldman, and Yixian Zheng. 2006. “A Mitotic Lamin B Matrix Induced by RanGTP Required for Spindle Assembly.” Science 311 (5769): 1887–93. https://doi.org/10.1126/science.1122771.

Vitre, Benjamin, Nikita Gudimchuk, Ranier Borda, Yumi Kim, John E Heuser, Don W Cleveland, and Ekaterina L Grishchuk. 2014. “Kinetochore-Microtubule Attachment throughout Mitosis Potentiated by the Elongated Stalk of the Kinetochore Kinesin CENP-E.” Molecular Biology of the Cell, 1–26. https://doi.org/10.1091/mbc.E14-01-0698.

Wang, Songyu, Fabian B. Romano, Christine M. Field, Tim J. Mitchison, and Tom A. Rapoport. 2013. “Multiple Mechanisms Determine ER Network Morphology during the Cell Cycle in Xenopus Egg Extracts.” Journal of Cell Biology 203 (5): 801–14. https://doi.org/10.1083/jcb.201308001.

Wang, Songyu, Hanna Tukachinsky, Fabian B. Romano, and Tom A. Rapoport. 2016. “Cooperation of the ER-Shaping Proteins Atlastin, Lunapark, and Reticulons to Generate a Tubular Membrane Network.” ELife 5 (September): 1–29. https://doi.org/10.7554/eLife.18605.

